# Rapid, activity-dependent intrinsic plasticity in the developing zebra finch auditory cortex

**DOI:** 10.1101/2023.02.07.527481

**Authors:** Yao Lu, Francesca Sciaccotta, Leah Kiely, Benjamin Bellanger, Alev Erisir, C Daniel Meliza

## Abstract

The acoustic environment an animal experiences early in life shapes the structure and function of its auditory system. This process of experience-dependent development is thought to be primarily orchestrated by potentiation and depression of synapses, but plasticity of intrinsic voltage dynamics may also contribute. Here we show that at the peak of the critical period for song memorization, neurons in the zebra finch caudal mesopallium, a cortical-level auditory area, can rapidly change their firing dynamics. This plasticity was only observed in birds that were reared in a complex acoustic and social environment, which also caused increased expression of the low-threshold potassium channel Kv1.1 in the plasma membrane and endoplasmic reticulum. Intrinsic plasticity depended on activity, was reversed by blocking low-threshold potassium currents, and was prevented by blocking intracellular calcium signaling. Taken together, these results suggest that Kv1.1 is rapidly mobilized to the plasma membrane by activity-dependent elevation of intracellular calcium. This produces a shift in the excitability and temporal integration of CM neurons that may be permissive for auditory learning in complex acoustic environments during a crucial period for the development of vocal perception and production.

## Introduction

The voltage-gated ion channels an individual neuron contains in its plasma membrane determine how it integrates synaptic inputs over time to produce patterns of action potentials, affecting its functional response properties (Llinás, 1988; Storm, 1988). The diverse intrinsic dynamics found within many brain areas contribute to sensory coding (Svirskis et al., 2002; Rothman and Manis, 2003; Padmanabhan and Urban, 2010), memory consolidation (Titley et al., 2017; Lisman et al., 2018), and maintaining network stability (Cudmore et al., 2010). Although the complement of channels expressed by individual neurons may primarily depend on the genetic programs that define cell types (Bishop et al., 2015), intrinsic dynamics often remain plastic, changing in response to activity and experience (Zhang and Linden, 2003; Daoudal and Debanne, 2003; McKay et al., 2009; Debanne et al., 2019; Santin and Schulz, 2019). Alongside synaptic plasticity, intrinsic plasticity may contribute substantially to experience-dependent functional changes during learning and development (Titley et al., 2017; Lisman et al., 2018; Daou and Margoliash, 2021), but in contrast to synaptic plasticity, the underlying mechanisms remain poorly understood.

The zebra finch (*Taeniopygia guttata*) is a social songbird that communicates with learned and innate vocalizations (Elie and Theunissen, 2016). As in many other species and sensory systems, development of the zebra finch auditory system is shaped by the acoustic environment experienced early in life (Amin et al., 2007; Woolley, 2012; Moore and Woolley, 2019). The intrinsic dynamics of neurons in the caudal mesopallium (CM), a cortical-level auditory area implicated in discriminating and learning species-specific vocalizations (Gentner and Margoliash, 2003; Meliza and Margoliash, 2012), are sensitive to acoustic experience during the developmental window when young males begin to memorize a song to copy (Chen and Meliza, 2020). Just after zebra finches fledge from the nest, almost all of the putative excitatory neurons in CM exhibit tonic dynamics, spiking continuously when depolarized. If the birds are reared in a breeding colony with many other families, within two weeks about half of these cells are phasic, spiking only at the onset of stimulation. If the birds are instead reared in abnormally quiet conditions, phasic neurons do not appear until much later in life, suggesting that intrinsic plasticity is an adaptive mechanism that permits normal auditory development in challenging acoustic conditions.

Phasic dynamics in CM depend on a low-threshold outward current (Chen and Meliza, 2018), and the experience-dependent emergence of phasic dynamics is correlated with expression of Kv1.1 (Chen and Meliza, 2020), a low-threshold voltage-gated potassium channel that facilitates phasic firing and is regulated by activity in a number of other systems (Brew and Forsythe, 1995; Bal and Oertel, 2001; Lu et al., 2004; Watanabe et al., 2014; Dehorter et al., 2015; Akter et al., 2018; Morgan et al., 2019). However, expression alone is not sufficient to cause a shift in intrinsic dynamics, because Kv1.1 must be exported to the plasma membrane, a process that may require activity-dependent elevation of intracellular calcium levels (Manganas et al., 2001; Rangaraju et al., 2010; Vacher and Trimmer, 2012; Galea et al., 2014). Indeed, at light microscopy resolution, much of the Kv1.1 staining in CM neurons is confined within the cell body, and in quiet-reared birds there are many more neurons that express Kv1.1 than exhibit phasic dynamics (Chen and Meliza, 2020). Thus, there may be a pool of ready-to-use Kv1.1 that could be trafficked to the plasma membrane in response to activity.

In the present study, we tested this model using several approaches. We examined how acoustic experience influences the expression and localization of Kv1.1 using confocal and electron microcopy in CM neurons from birds between 30–35 days of age, confirming the existence of an experience-dependent pool of Kv1.1 in the endoplasmic reticulum (ER) that could be exported to the membrane. We made whole-cell recordings from putative excitatory neurons during this window to test the prediction that individual phasic neurons would all express Kv1.1 but that the converse would not be the case, because neurons with high levels of Kv1.1 in the ER might not have enough in the plasma membrane to cause phasic spiking. Finally, we tested whether colony-reared neurons, which have high levels of Kv1.1 expression, would become more phasic when repeatedly stimulated to spike, using pharmacology to confirm that this change in intrinsic dynamics depends on low-threshold potassium currents.

## Materials and Methods

### Animals

All procedures were performed according to National Institutes of Health guidelines and protocols approved by the University of Virginia Institutional Animal Care and Use committee. Experiments used slices from juvenile zebra finches (*Taeniopygia guttata*; 52 birds) between 30–35 days after hatch. Finches were bred in our colony from 25 different pairs. All birds received finch seed (Abba Products, Hillside, NJ) and water ad libitum and were kept on a 16:8 h light-dark schedule in temperature- and humidity-controlled rooms (22–24 °C).

### Experimental rearing conditions

All of the animals in the study were bred in individual cages that were initially placed in a room housing dozens of male and female finches of varying ages. Colony-reared (CR) chicks remained with their parents in the colony room until they were used in an experiment. Pair-reared (PR) chicks were moved with their parents into an acoustic isolation box (Eckel Industries) within 7 d after the first egg hatched.

### Immunofluorescence

We compared Kv1.1 expression in CR and PR animals using a similar procedure to the one described previously (Chen and Meliza, 2020). Animals were administered a lethal intramuscular injection of Euthasol and perfused transcardially with a 10 U/mL solution of sodium heparin in PBS (in mM: 10 Na2HPO4, 154 NaCl, pH 7.4) followed by 4% paraformaldehyde (in PBS). Brains were immediately removed from the skull, postfixed overnight in 4% paraformaldehyde at 4 °C, and then transferred to PBS. Brains were blocked along the same plane used for making acute slices, and sections were cut at 60 µm on a vibratome and stored in PBS with 0.05% sodium azide until ready for staining. Sections were rinsed four times in PBS and then incubated in citrate buffer (0.01 M, pH 8.5) at 80 °C for 25 minutes. The sections were cooled to room temperature, washed twice in PBS, permeabilized in PBS-T (0.1% Tween in PBS), and blocked in 5% goat serum and 2% glycine in PBS-T. The tissue was stained for Kv1.1 using a monoclonal mouse antibody (1:500 in blocking solution; UC Davis/NIH Neuromab clone K20/78; RRID:AB_10673165) for 60–72 h at 4 °C. Sections were washed 4 times for 15 min in PBS-T and then incubated with a fluorescent secondary antibody (1:2000 in PBS-T; goat anti-mouse IgG1 conjugated to Alexa Fluor 488; Invitrogen; RRID:AB_2534069). The tissue was washed twice with PBS, counterstained for Nissl (Neurotrace 640/660, 1:1000 in PBS; ThermoFisher, catalog N21483; RRID:AB_2572212), washed four times with PBS, and then mounted and coverslipped in Prolong Gold with DAPI (ThermoFisher, catalog P36934; RRID:SCR_015961). Stained tissue was imaged on a Zeiss LSM 800 confocal microscope using a 40X objective (water immersion, NA 1.2) in stacks with 0.44 µm between optical sections. At least three stacks were imaged in each section, and two sections were imaged per animal for a total of 6 sections. To quantify expression, we counted the number of Kv1.1-positive neurons in each stack manually in Imaris (version 9.2.1) after using the automatic background subtraction, with the default smoothing radius of 39.9 µm. The total number of neurons in each stack was counted from the DAPI channel using the automatic spot detection function in Imaris, followed by a manual check. Samples were processed and analyzed in two batches, and each batch contained samples from both conditions. Identical laser power and gain settings were used for the Kv1.1 channel in all the samples from each batch, but the DAPI settings were adjusted freely to optimize signal quality. Experimenters were blind to animal identity and rearing condition throughout staining, image collection, and analysis.

Slices used for acute recordings were stained for Kv1.1 and biocytin using a similar procedure with modifications for thick sections. Permeabilization, blocking, and staining solutions were prepared with a stronger detergent (0.3% Triton-X instead of 0.1% Tween). The blocking step was hours; the primary antibody incubation was 7 d; and the secondary staining incubation lasted for 20 h and included streptavidin-conjugated Alexa Fluor 635 (1:1000, Molecular Probes) to label biocytin. The biocytin-filled neuron was imaged with the same microscope, objective, and spacing between optical sections described above.

Cells were only included for analysis if the recorded neuron could be unambiguously identified in the slice (sometimes biocytin expelled from the pipette during patching was taken up by neighboring cells) and if Kv1.1 staining was good enough to clearly see positive neurons at the depth of the cell within the section. Included neurons were classified as Kv1.1-negative or Kv1.1-positive by an observer (C.D.M.) who was blind to the electrophysiological characteristics of the cell. Cells were deemed to be positive only if they had clearly defined, punctate Kv1.1 staining that was elevated higher than the background, co-extensive with the biocytin signal in the soma, and showing a region of lower staining corresponding to the nucleus (e.g., Fig. 1A). Almost all (*n* = 24/27) of the filled neurons were positive by these criteria. This proportion is much higher than the roughly 50% we reported previously (Chen and Meliza, 2020). Differences in staining technique (these sections were much thicker and were incubated in primary antibody much longer) and selection criteria (this sample only includes neurons that we were able to successfully patch) may explain this discrepancy.

**Fig. 1.**
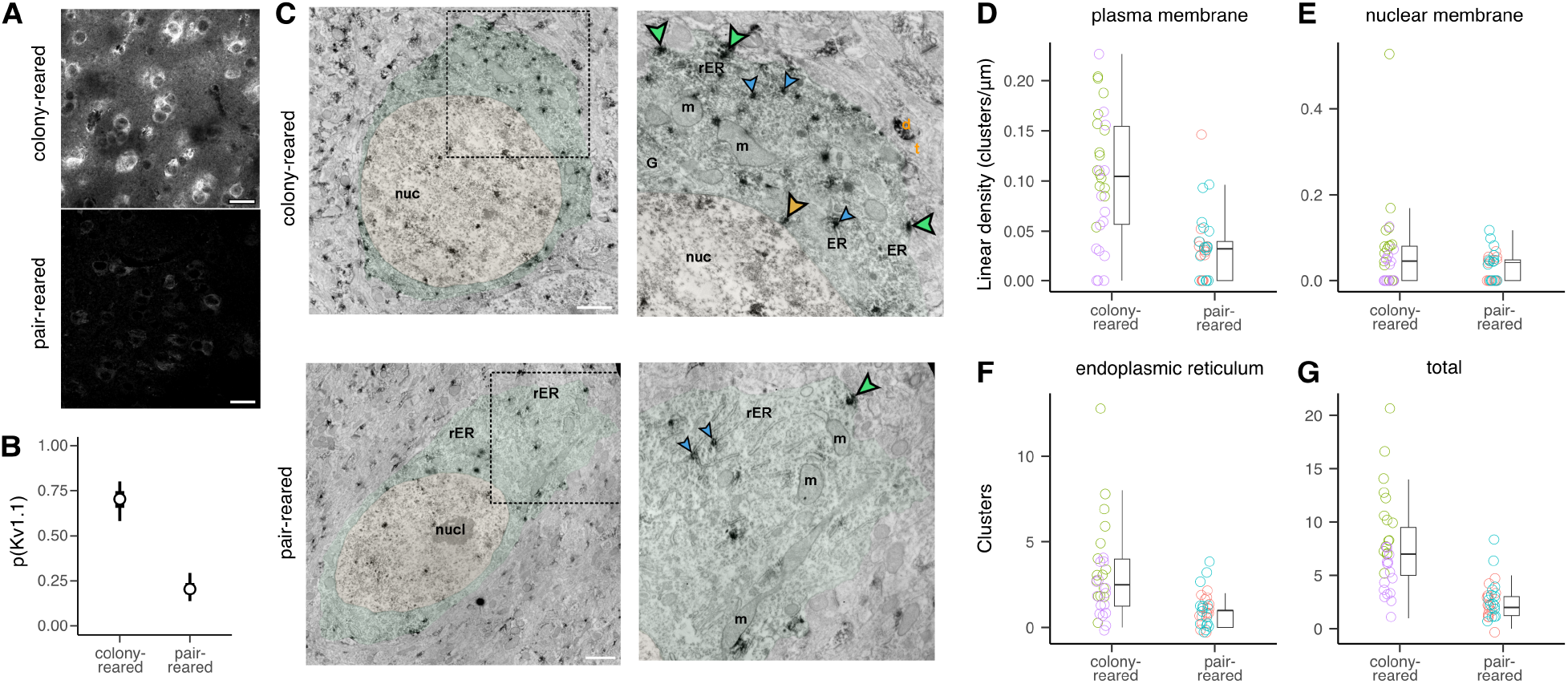
Early auditory experience modulates expression and localization of Kv1.1. (**A**) Examples of *α*-Kv1.1 staining in CM at P30–P35 in CR and PR zebra finches. Images are single confocal slices (0.44 µm) with identical laser, gain, and contrast settings. Scale bar: 20 µm. (**B**) Proportion of Kv1.1-positive neurons in each rearing condition from *n* = 2 CR and *n* = 3 PR chicks. Hollow circles show means for each condition; thick and thin whiskers show 50% and 90% credible intervals for the mean (GLMM; see materials and methods). There are many more immunopositive neurons in CR birds (GLMM: log-odds ratio = 2.2*±* 0.36, *p <* 0.001). (**C**) Composite electron micrograph of a Kv1.1-immunopositive cell from a CR bird (top) and a PR bird (bottom). The nucleus (yellow) and cytoplasm (green) are pseudocolored. Scale bar: 1 µm. The right panel shows detail of the area indicated by a dashed line in the left panel. DAB staining product accumulated in discrete clusters on the plasma membrane (green arrowheads), the endoplasmic reticulum (blue arrowheads) and the cytosolic side of the nuclear membrane (orange arrowhead). Abbreviations: m, mitochondria; ER, endoplasmic reticulum; rER, rough ER; G, Golgi apparatus; nuc, nucleus; nucl, nucleolus; d, dendrite; t, terminal. (**D**) Comparison of the linear density (DAB clusters per µm of membrane) of Kv1.1 staining on the plasma membrane for CR and PR conditions. Hollow circles correspond to individual neurons, with color indicating the bird (*n* = 15 neurons from 2 birds in each condition). Boxplots indicate median (horizontal line), interquartile range (central box) and the largest value no further than 1.5 times the interquartile range (whiskers). The density of Kv1.1 on the plasma membrane is higher in CR birds (Wilcoxon test: *W* = 739.5, *p <* 0.001).

The biocytin-filled neurons were also analyzed using an automated pipeline to quantify expression by the number of puncta in the soma. The 3D stacks were first cropped in Imaris to include only the volume immediately surrounding the soma. One cell was discarded because of a blood vessel within this volume that stained strongly in the Kv1.1 channel. The cropped image was exported for analysis in Cell Profiler (version 4.2.4). Illumination was corrected separately in the biocytin and Kv1.1 channels using the regular illumination function with a polynomial smoothing function. An Otsu global method strategy with three-class thresholding to identify objects from 6–14 pixels in diameter in the Kv1.1 channel. The Otsu global method strategy with two-class thresholding was used to identify the soma from the biocytin channel. Finally, the ClassifyObjects method was used to count Kv1.1 puncta located with the cell soma. Density was calculated as the total number of colocated puncta divided by the sum of the area in each optical section times the section thickness (0.44 µm).

### Immuno-electron microscopy

Animals were deeply anesthetized with an overdose of euthasol and transcardially perfused with Tyrode’s solution followed by 100 mL of fixative solution containing 4% paraformaldehyde and 0.1– 0.3% glutaraldehyde in PBS. Brains were immediately removed from the skull, postfixed overnight in 4% paraformaldehyde at 4°C, blocked for modified coronal sections, and cut at 60 µm on a vibratome. Sections were rinsed in 1% sodium borohydride and stored in 0.05% sodium azide in PBS at 4°C.

Kv1.1 was stained as for immunofluorescence, with the following changes: sections were incubated in citrate buffer (0.01 M, pH 8.5) at 80°C for 2 × 15 min. Sections were cooled to room temperature, washed three times in PBS, and pre-incubated in a solution with 1% bovine serum albumin (BSA) and 0.1% Triton-X in PBS for only 30 min. Sections were stained with a solution of primary antibody (1:500), 1% BSA, and 0.05% sodium azide for 72 h at 4°C on a shaker. The secondary antibody incubation was with horse anti-mouse IgG conjugated to biotin (1:100 in PBS; Vector Labs, catalog BA-2000-1.5) for 2 h at room temperature. Sections were treated with avidin-biotinylated peroxidase (VECTASTAIN ABC-HRP staining kit; Vector Labs, catalog PK-4000) for 2 h and developed in a solution of hydrogen peroxide (H2O2) and 0.05% diaminobenzidine (DAB) for 2–7 min.

Sections were embedded for electron microscopy by incubation with 1% osmium tetroxide (OsO4) in 0.1 M sodium phosphate buffer (PB, pH 7.4) for 1 h then with filtered 4% uranyl acetate in 70% alcohol for 1 h, followed by dehydration in acetone and treatment with a 1:1 acetone/resin mixture overnight. The following day, sections were transferred to full resin and left overnight. Sections were then flat-embedded between two aclar sheets and cured in a 60°C oven overnight. The region of interest from each section was placed in a BEEM capsule (EMS, Hatfield, PA). The capsules were filled with resin and cured at 60°C for 24–48 h, or until polymerized. The region of interest (now capsule-embedded) was traced with a camera lucida and trimmed down to a 1mm × 2mm trapezoid. Ultrathin sections of 50-80nm thickness were collected on 400 mesh copper grids (Ted Pella, Redding, CA) using an ultramicrotome (Ultracut UCT7; Leica, Buffalo Grove, IL). The ultrathin sections were examined on a JEOL1010 electron microscope equipped with a 16-megapixel CCD camera (SIA). Images for quantitative and immuno-labeled terminal analysis were taken at 8,000–15,000X magnification, yielding a resolution of 576.06–1134.92 pixels/µm. Kv1.1 localization was analyzed in 15 neurons from each bird (*n* = 2 birds per condition) by counting the number of DAB clusters associated with the endoplasmic reticulum, the nuclear membrane, and the plasma membrane. Clusters that were not clearly associated with one of these compartments were excluded because the DAB might have diffused from a structure in an adjacent section. The plasma membrane and nuclear membrane counts were normalized relative to the length of the membrane in the section. Counts were pooled from the two birds within each condition and compared between conditions using Mann-Whitney U tests.

### Brain slice preparation and electrophysiology

Whole-cell recordings were obtained from acute slices of the caudal mesopallium. Birds were administered a lethal intramuscular injection of Euthasol (pentobarbital sodium and phenytoin sodium; 200 mg/kg; Hospira) and perfused transcardially with ice-cold cutting buffer (in mM: 92 NaCl, 2.5 KCl, 1.2 NaH2PO4, 30 NaHCO3, 20 HEPES, 25 glucose, 5 sodium ascorbate, 2 thiourea, 3 sodium pyruvate, 10 MgSO4·7H2O, 0.5 CaCl2·2H2O, pH 7.3–7.4, 310–330 mmol/kg). The brain was blocked using a custom 3D printed brain holder (http://3dprint.nih.gov/discover/3dpx-003953) on a dorsorostral to ventrocaudal plane pitched 65 degrees forward of the transverse plane, so that slices through the auditory pallium would be approximately orthogonal to the ventral mesopallial lamina (LMV) (Chen and Meliza, 2018). Sections 300 µm thick were cut in room-temperature cutting buffer on a VF-200 or VF-500 Compresstome (Precisionary Instruments), and then transferred to 32°C cutting buffer for 10–15 min for initial recovery. Slices were then transferred to room-temperature holding buffer (in mM: 92 NaCl, 2.5 KCl, 1.2 NaH2PO4, 30 NaHCO3, 20 HEPES, 25 glucose, 5 sodium ascorbate, 2 thiourea, 3 sodium pyruvate, 2 MgSO4·7H2O, 2 CaCl2·2H2O, pH 7.3–7.4, 300–310 mmol/kg) to recover for at least 1 h before use in recordings. All solutions were bubbled continuously with 95% O2·5% CO2 mixture starting at least 10 min before use.

Slice recordings were conducted in a RC-26G recording chamber (Warner Instruments) perfused with standard recording ACSF (in mM: 124 NaCl, 2.5 KCl, 1.2 NaH2PO4, 24 NaHCO3, 5 HEPES, 12.5 glucose, 2 MgSO4·7H2O, 2 CaCl2·2H2O; pH 7.3–7.4, 300–310 mmol/kg) at a rate of 1–2 mL/min at 32°C. Whole-cell patch-clamp recordings were obtained under 40X infrared (900 nm) DIC optics. CM was located relative to the lamina mesopallialis ventralis (LMV) and the internal occipital capsule (CIO), which both comprise dense myelinated fibers visible as dark bands under bright-field or IR illumination. Recording pipettes were pulled from filamented borosilicate glass pipettes (1.5 mm outer diameter, 1.10 mm inner diameter; Sutter Instruments) using a P-1000 Micropipette Puller (Sutter Instruments) and were filled with internal solution (in mM: 135 K-gluconate, 10 HEPES, 8 NaCl, 0.1 EGTA, 4 MgATP, 0.3 NaGTP, 10 Na-phosphocreatine, pH 7.2–7.3, 300–310 mmol/kg). In some recordings, 0.2% biocytin (Sigma) was included in the internal solution.

Electrodes had a resistance of 4–8 MΩ in the bath. Voltages were amplified with a Multiclamp 700B amplifier (Molecular Devices) in current-clamp mode, low-pass filtered at 10 kHz, and digitized at 50 kHz with a Digidata 1440A. Pipette capacitance was neutralized, and 8–12 MΩ of series resistance was subtracted by bridge balance. Recorded voltage was corrected offline for a liquid junction potential of 11.6 mV (as measured at 32°C). Current injections and data collection were controlled by pCLAMP (version 10.4; Molecular Devices). Neurons were excluded from analysis if the resting membrane potential was above –55 mV or if action potentials failed to cross –10 mV.

After a brief test to ascertain how much current was needed to evoke spikes, each neuron was recorded in epochs of 20 sweeps. Each sweep comprised a 2 s depolarizing step current and two 500 ms hyperpolarizing steps. The amplitude of the depolarizing pulse was systematically varied within each epoch over a range that was set so that at least 10 sweeps in each epoch evoked action potentials without eliciting depolarization block. The range of current steps was adjusted throughout the recording as needed. For the colocalization experiments, the recordings were terminated after one or two epochs by slowly withdrawing the electrode from the cell until it detached and formed an outside-out patch, as evidenced by a sharp increase in resistance. The slice was allowed to rest in the recording chamber for 5–10 min and then fixed in 4% formaldehyde for 15 h prior to staining. In plasticity experiments, the standard condition was to record epochs one after another as long as the cell appeared to remain healthy. Additional exclusion criteria were applied *post hoc* (see below). In the *reduced stimulation* condition, after recording a single epoch at the beginning of the experiment, the cell was voltage-clamped at resting potential for at least 400 s, followed by another recording epoch. During the voltage-clamp interval, performing a seal test (–2 mV pulses at 100 Hz) tended to help avoid resealing.

### Pharmacology

To test whether blocking potassium currents would reverse intrinsic plasticity, recordings were performed in standard ACSF with an added cocktail of drugs to block fast synaptic transmission: 10 *µ*M (*±*)-3-(2-carboxyp-iperazin-4-yl)propyl-1-phosphonic acid (CPP; Alomone Labs), 20 *µ*M 6-cyano-7-nitroquinoxaline-2,3-dione disodium salt hydrate (CNQX; Alomone Labs), and 10 *µ*M (+)-bicuculline (Sigma). After the neuron became phasic (4–11 epochs), 2 mM 4-aminopyridine (4-AP; Sigma) was added to the perfusate. In experiments to reduce intracellular Ca^2+^ activity, slices were incubated in ACSF containing membrane-permeant AM variant of the fast exogenous Ca^2+^ chelator 1,2-Bis(2-aminophenoxy)ethane-N,N,N’,N’-tetraacetic acid tetrakis(acetoxymethyl ester) (BAPTA-AM, 10 *µ*M, Sigma-Aldrich) for 30 min and then returned to normal ACSF for another 30 min before recording. Attempts to prepare higher concentrations of BAPTA-AM resulted in precipitate formation.

### Signal processing and analysis

Recordings were analyzed using custom Python code (available upon publication at https://github.com/melizalab/cm_plasticity). If the average series resistance deviated in any epoch by more than 30% from baseline, the resting membrane potential increased by more than 10 mV, or the cell spiked spontaneously more than five times, that epoch and all the ones that followed were excluded. Spontaneous firing was used as an exclusion criterion because it indicated poor health, it interfered with analysis, and it introduced uncontrolled variation in activity. Input resistance was not used as an exclusion criterion because it was expected to change with an increase or decrease in low-threshold potassium conductance. After excluding bad epochs, individual sweeps were excluded if the resting membrane potential was transiently unstable, defined as a fluctuation more than 10 times the median absolute deviance. Action potentials were detected following previously described procedures (Chen and Meliza, 2018, 2020). The response duration in each sweep was measured as the time elapsed between the first and the last spikes of the depolarizing pulse. In responses with only one spike, the duration was defined as the width of the spike plus the duration of the afterhyperpolarization. The rheobase was defined as the average of the largest current that produced no spikes and the the smallest current that produced one or more spikes. The slope of the frequency-current (*f-I*) curve was calculated for each epoch from the above-threshold sweeps as the average of the change in firing rate divided by the average of the increase in current. Current-voltage (*I-V*) curves were measured from all sweeps using the steady-state voltages produced by both depolarizing and hyperpolarizing pulses. For the depolarizing currents, the voltage values were calculated from the last 500 ms of the pulse and were only included if there were no action potentials during this interval.

### Experimental Design and Statistical Analysis

Intrinsic spiking dynamics were quantified using two measures: the average duration of spiking in response to depolarizing step currents, and the average slope of the f-I curve. In contrast to our previous study, we did not log-transform the durations, because the residuals of statistical models using untransformed durations (i.e., arithmetic difference) were closer to a normal distribution than the residuals in models with log-transformed durations (i.e., geometric difference), as assessed by visual examination of a Q-Q plot and the Shapiro-Wilk test. Duration and slope were either compared to the expression of Kv1.1 measured with immunohistochemistry or tracked over time to determine under what conditions intrinsic dynamics were stable or plastic. At least 7 animals were used for slice experiments in each condition tested; this sample size was chosen based on previously published studies from our lab examining intrinsic dynamics in these neurons (Chen and Meliza, 2018, 2020). The experimenter was aware of the rearing conditions of the animals and the experimental conditions during recording. The experimental design was not preregistered.

Statistical inference was performed for confocal and electrophysiology data using mixed-effects models with bird and family identity as random effects to account for repeated measures. When the dependent measure was spike duration, individual sweeps were treated as the unit of observation, and a random intercept (and slope, if applicable) was added for each neuron. In the experiments measuring intrinsic plasticity, the four experimental conditions were analyzed in a single model. Fixed-effect contrasts were planned to compare the last good epoch with the first epoch, and the experimental conditions (PR birds, BAPTA-AM, and reduced stimulation) with the standard condition (CR birds, standard ACSF, continuous stimulation). The reversal experiment was analyzed separately. Only tonic neurons that became phasic over the course of recording were included, and the epoch after applying 4-AP was compared to the epoch before it was applied. The proportion of cells expressing Kv1.1 in CR and PR birds was estimated using a mixed-effects model with a binomial error distribution. Image stacks were treated as the unit of observation, with section, bird, and staining batch as random effects, and rearing condition as the sole fixed effect. All of the models were fit in R using the *lme4* package (version 1.1-29), and the effects, post-hoc comparisons, confidence intervals, and p values were calculated using the *emmeans* package (version 1.8.1). Satterthwaite approximations were used to estimate effective degrees of freedom, which are denoted in the text as subscripts and reflect both the sample size and within-group variances; thus, not all effects tested in a model have the same degrees of freedom.

Non-parametric statistical tests (Wilcoxon rank-sum test) were used for the immuno-EM data because of the small number of cells and animals.

## Results

### Kv1.1 expression and localization depend on experience

To test whether there is an experience-dependent pool of ready-to-use Kv1.1 that could support rapid changes of intrinsic dynamics, we used immunohistochemistry to compare expression and localization of Kv1.1 in CM neurons from colony-reared (CR) and pair-reared (PR) zebra finch chicks between P30–P35 (days post hatch). In confocal images, CR birds have higher levels of Kv1.1 expression, with a much larger proportion of neurons expressing detectable levels of Kv1.1 (Fig. 1A,B). This result is consistent with our earlier finding that Kv1.1 expression is higher in CR birds compared to female-reared birds, who do not hear any song (Chen and Meliza, 2020), and it confirms that Kv1.1 expression occurs in response to a noisy, complex acoustic environment rather than exposure to a tutor song. We observed the same spatial pattern of expression as previously, with weak staining throughout the neuropil and intense, punctate immunoreactivity in the cell body around the nucleus.

To determine what subcellular compartments contained Kv1.1, we used electron microscopy to compare immunolabeled tissue from CR and PR birds. Kv1.1 protein was associated with the plasma membrane, the ER, and the cytosolic side of the nuclear membrane (Fig. 1C). A similar pattern is seen in the mammalian neocortex (Goldberg et al., 2008). Staining was observed in dendrites and spines near the postsynaptic density; these occurrences were not scored but may also represent Kv1.1 that could contribute to membrane voltage dynamics. Staining was also present around mitochondria and in regions of the cytoplasm not obviously occupied by any organelles. As in the confocal images, there was more Kv1.1 in the CR birds (Fig. 1G). CR birds had more dense staining along the plasma membrane (Fig. 1D) and more clusters in the ER (Fig. 1F), but there was no difference in staining of the nuclear membrane (Fig. 1E). The cytosolic area was statistically indistinguishable between CR and PR birds (Wilcoxon test: *W* = 466, *p* = 0.81), so the higher number of clusters found in the plasma membrane and ER of CR birds is likely to reflect increased levels of expression. Thus, birds that experienced a complex acoustic environment had both more Kv1.1 at the plasma membrane and a larger pool of Kv1.1 in the ER that could be exported to the plasma membrane.

The linear density of staining on the nuclear membrane for CR and PR birds, same format as **D**. The difference is not significant (*W* = 541.5, *p* = 0.16). (**F**) The number of clusters associated with the ER is greater in cells from CR birds compared to PR birds (*W* = 704.5, *p <* 0.001). (**G**) The total number of clusters associated with ER, plasma membrane, and nuclear membrane is higher in cells from CR birds compared to PR birds (*W* = 807, *p <* 0.001).

### Kv1.1 is not sufficient for phasic excitability

The existence of a ready-to-use pool of Kv1.1 implies that expression of Kv1.1 is necessary but not sufficient for phasic excitability. To test this prediction, we performed whole-cell recordings from neurons in acute slices of CM from colony-reared chicks. Cells were stimulated with 2 s depolarizing current steps at a range of amplitudes, and as we have seen previously, some neurons responded throughout the depolarization (tonic spiking) while others responded only near the onset (phasic spiking). This difference was quantified by averaging the duration of spiking across all the suprathreshold amplitudes. Longer durations correspond to tonic spiking (e.g., Fig. 2A,B), and shorter durations correspond to phasic spiking (e.g., Fig. 2C). The recording pipette was filled with biocytin, and after the recording was completed, the slices were stained for biotin and Kv1.1 to determine what, if any, relationship there was between intrinsic dynamics and Kv1.1 expression.

**Fig. 2.**
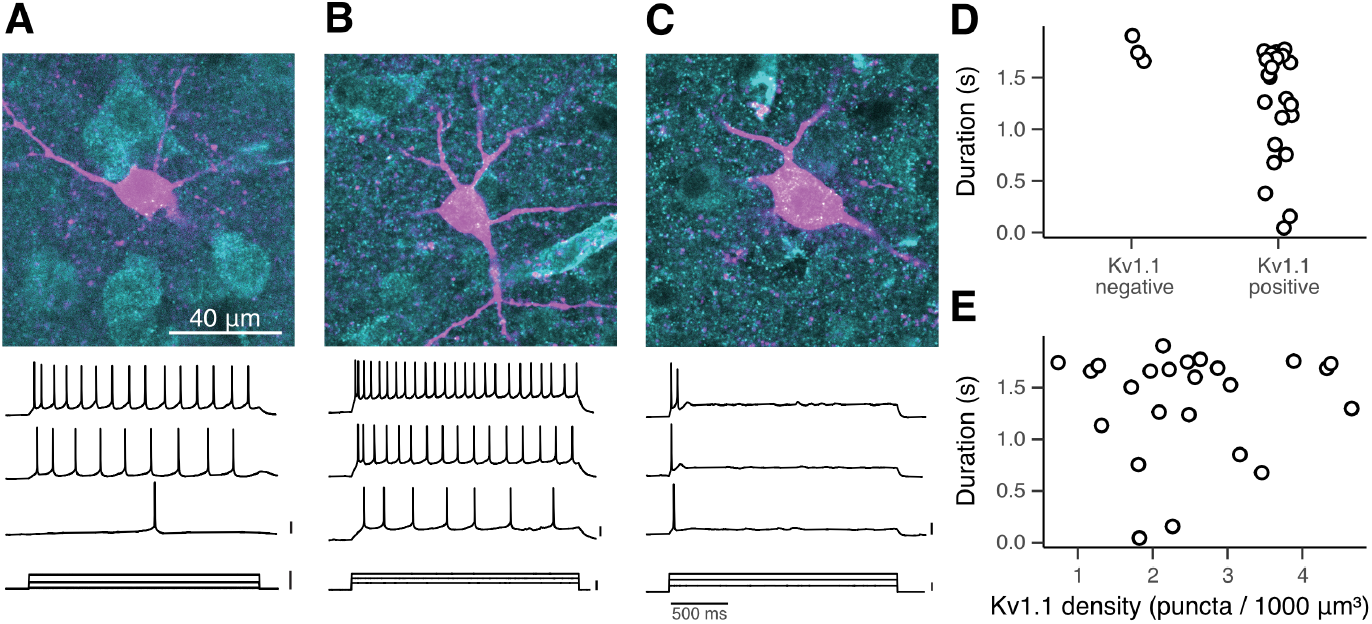
Kv1.1 is not sufficient for phasic excitability. (**A**) Example of a Kv1.1-negative tonic neuron. Image is a single confocal slice (0.44 µm) showing fluorescent staining for Kv1.1 (cyan) and biocytin (magenta). Scale bar: 40 µm. Bottom shows exemplar voltage traces in response to step current injections. Scale bars: 20 mV, 50 pA, 500 ms. (**B**) Example of a Kv1.1-positive tonic neuron. (**C**) Example of a Kv1.1-positive phasic neuron. (**D**) Average response duration for Kv1.1-negative (*n* = 3) and Kv1.1-positive (*n* = 24) neurons. Symbols represent single neurons and have been jittered horizontally for clarity. (**E**) Average response duration versus Kv1.1 staining, measured as the density of puncta detected within the stained soma. The correlation is not significant (Spearman correlation: *ρ*_22_ = 0.08, *p* = 0.70).

As illustrated in Figure 2A–C, biocytin-filled neurons had quasi-pyramidal somata with dendrites radiating in all directions. All but one of the neurons with good staining in the dendrites (*n* = 26/27) had spines and broad action potentials (mean width at half-height *±* SD: 1.9 *±* 0.51 ms), identifying them as putative excitatory neurons. The remaining neuron had varicose dendrites and fast, narrow spikes (0.89 ms at half-height), marking it as an inhibitory neuron, so we excluded it from further analysis. Only a small proportion (11%) of the excitatory, biocytin-filled neurons were negative for Kv1.1, and they were all tonic (Fig. 2A,D). In contrast, Kv1.1-positive neurons exhibited dynamics ranging from tonic (Fig. 2B) to highly phasic (Fig. 2C). Because we only recorded from a few Kv1.1-negative cells, it was not possible to make a meaningful statistical test of whether Kv1.1 expression was necessary for phasic firing, but it was clearly not sufficient, as there were many tonic neurons that expressed Kv1.1. Indeed, when we quantified Kv1.1 expression by the density of immunopositive puncta in the soma, there was no correlation with firing duration (Fig. 2E), and the neurons that stained the most strongly for Kv1.1 were tonic.

### Rapid, experience-dependent plasticity of intrinsic dynamics

If the ER and other intracellular locations contain a pool of Kv1.1, then neurons should be able to rapidly switch from tonic to phasic spiking by exporting Kv1.1 to the plasma membrane. To test this hypothesis, we tracked evoked spiking patterns for 100–2500 s in whole-cell recordings from CR chicks (Fig. 3A). Many neurons exhibited a shift during the recording from tonic to phasic spiking (Fig. 3B,C), along with an increase in the rheobase and a decrease in the slope of the relationship between firing rate and injected current (f-I curve; Fig. 3E,F). Phasic spiking was typically accompanied by the emergence of outward rectification at around –60 mV (Fig. 3D), which is characteristic of a low-threshold voltage-gated potassium current. Resting membrane voltage and input resistance tended to decrease or remain stable.

**Fig. 3.**
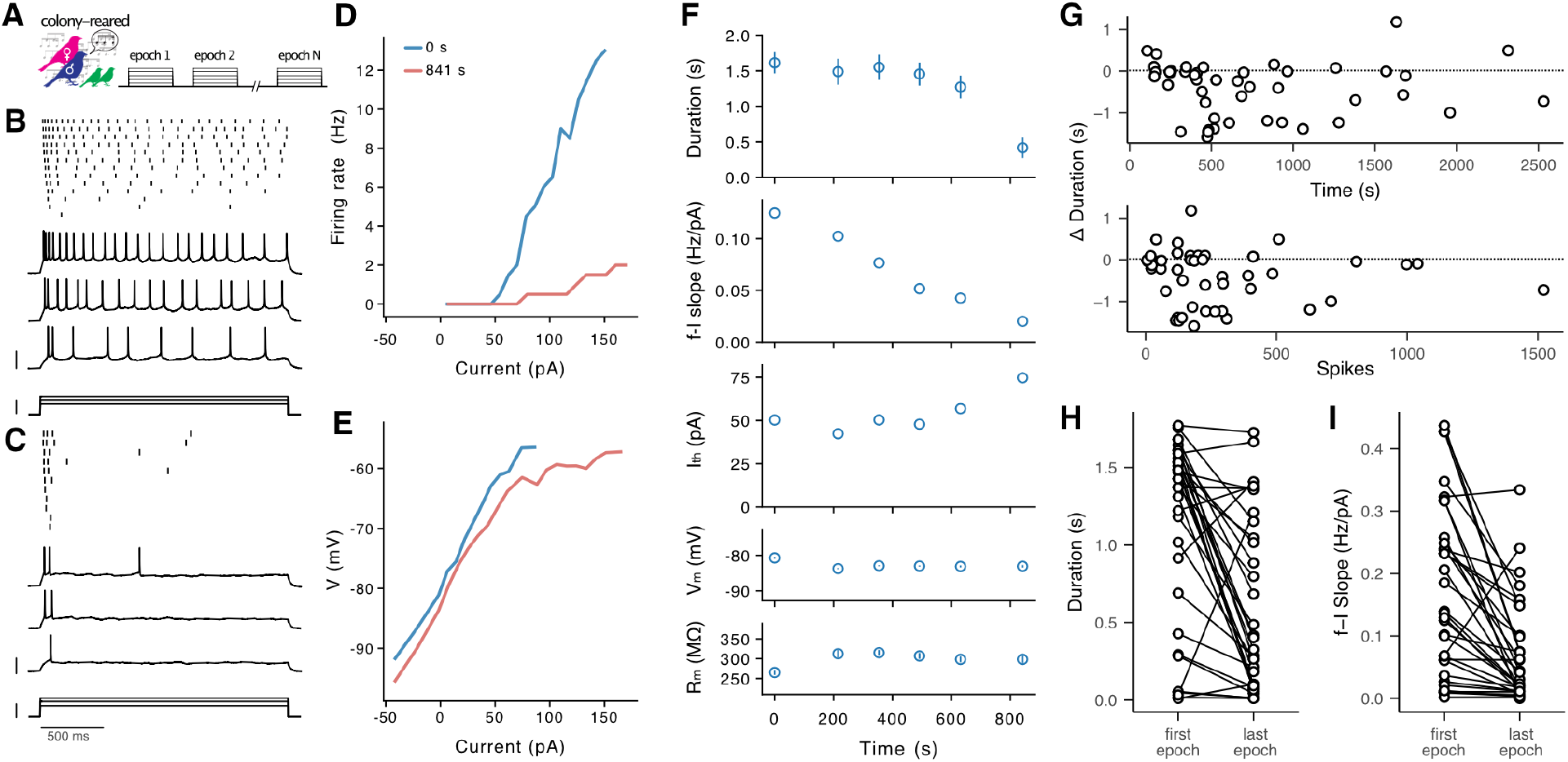
Rapid emergence of phasic excitability in colony-reared (CR) birds. (**A**) Neurons from colony-reared chicks were stimulated with epochs of depolarizing step currents 2 s in duration with varying amplitude. (**B**) Tonic responses of an exemplar neuron immediately after establishing the whole-cell recording. Top is a raster plot showing spike times. Below are voltage and current traces for three selected trials. Scale bars: 40 mV, 100 pA, 500 ms. (**C**) Phasic response of the same exemplar neuron 841 s after the start of the recording. Same format as in (A). (**D**) Firing rate versus injected current amplitude (f-I curve) for the first and last epochs. (**E**) Steady-state voltage versus injected current amplitude for the first and last epochs of the recording. The line is plotted only when the cell did not spike during the measurement window. Note the emergence of strong outward rectification in the latter epoch. Firing duration, slope of frequency vs. current, rheobase (Ith), resting potential (Vm), and resting input resistance (Rm) for each epoch in the recording. Vertical lines show standard error of mean for duration, Vm, and Rm, which were averaged across all the sweeps in the epoch. Pauses in the recording occurred when the range of stimulating currents was adjusted to ensure an adequate number of traces included spikes (see Methods). (**G**) Change in duration from first to last epoch versus total recording time (top) and total number of spikes (bottom). Each symbol corresponds to the last epoch recorded in an individual neuron. (**H**) Comparison of average firing duration in the first and last epochs for *n* = 33 neurons in colony-reared birds where the recording lasted at least 400 s. The average change was *−*0.55 *±* 0.09 s (LMM: *t*_74.7_ = *−*6.0, *p <* 0.001). (**I**) Comparison of frequency-current slope in first and last epochs for the same neurons. The average change was*−* 0.100 *±*0.018 Hz/pA (*t*_76_ = *−*5.52, *p <* 0.001).

Large changes in intrinsic dynamics were observed only after about 400 s of recording, and persisted throughout recordings that lasted for over 30 min (Fig. 3G). Not every neuron was plastic. When intrinsic dynamics changed, it was almost always in the direction of more phasic responses. Nearly half of the neurons (43%, *n* = 10/23) decreased average spiking duration by over 1 s, whereas only a single highly phasic neuron became tonic. On average, the shift was toward more phasic spiking (Fig. 3H) and lower excitability (Fig. 3I). There was no statistically significant difference between males and females in either the initial duration (LMM: *β* = 0.001 s, *F*_1,8.8_ = 0.004, *p* = 0.95) or the final duration (*β* = 0.0006 s, *F*_1,28_ = 0.002, *p* = 0.96).

If this shift in intrinsic dynamics is caused by export of Kv1.1 to the plasma membrane, then it should only occur in animals that have a readily available pool of intracellular Kv1.1. Pair-reared birds have lower levels of Kv1.1 expression in the ER and overall (Fig. 1), so we expected that the intrinsic dynamics in neurons from PR birds would be much less plastic compared to what we observed in CR birds. This was indeed the case. As illustrated in Figure 4, although there was a tendency for neurons in PR birds to become less excitable, their firing patterns remained tonic. Decreased excitability was sometimes accompanied by a shift in the I-V curve at hyperpolarized voltages (Fig. 4E), but the low-threshold outward current that appeared in plastic CR neurons (Fig. 3E) was not observed. Overall, in PR birds there was no significant change in firing duration (Fig. 4H), even in recordings lasting close to 30 min (Fig. 4G). There was a modest but significant decrease in f-I slope (Fig. 4I), suggesting that other mechanisms may contribute to the decrease in excitability.

**Fig. 4.**
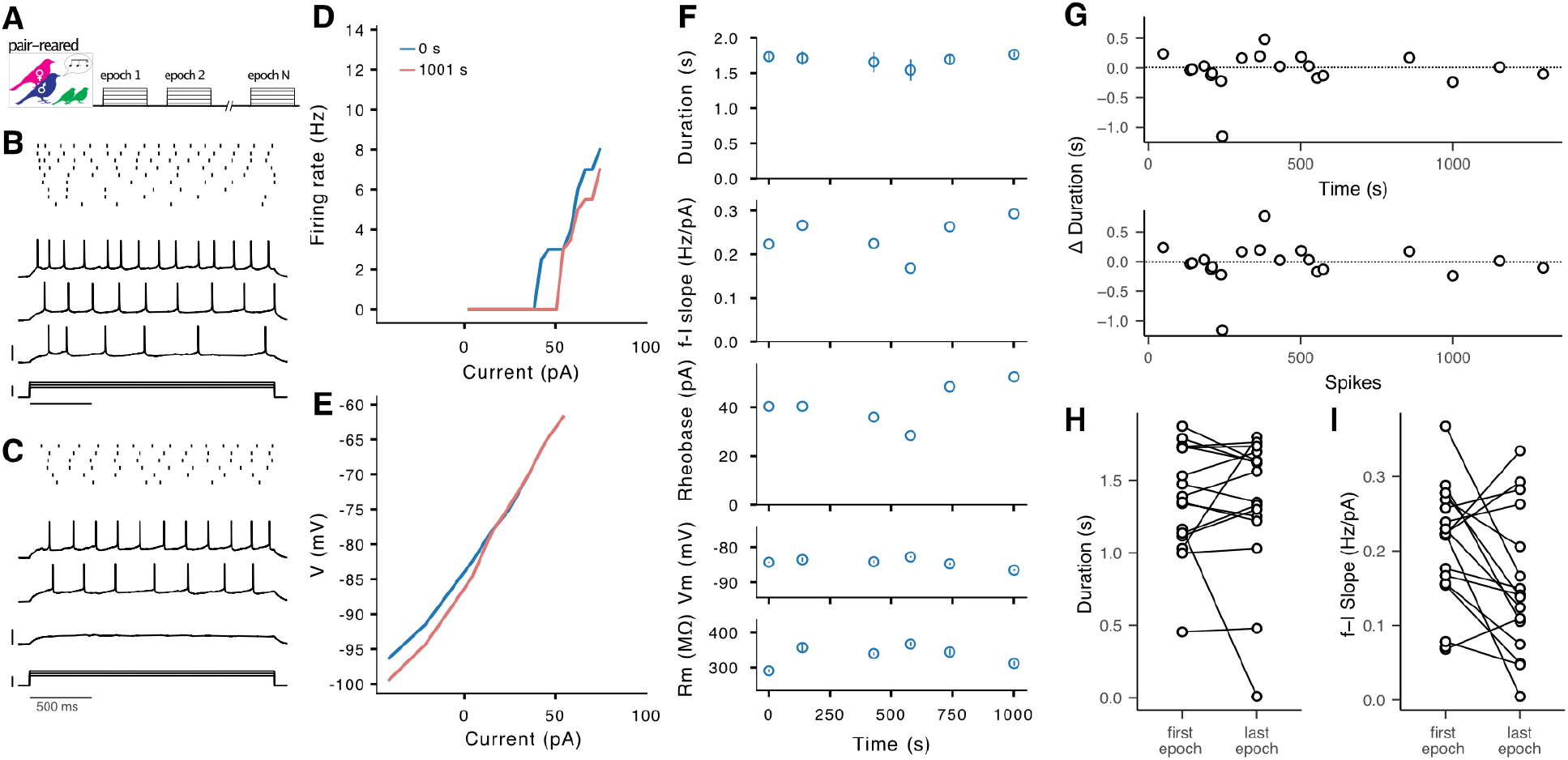
Tonic dynamics are stable in pair-reared (PR) birds. (**A**) Experimental design: neurons from PR chicks were stimulated with epochs of depolarizing step currents. (**B–F**) Responses of an exemplar neuron to depolarizing step currents immediately after establishing the whole-cell recording and 1001 s later. Same format as Figure 3B–F. (**G**) Change in duration from first to last epoch versus total recording time (top) and total number of spikes (bottom). (**H**) Comparison of average firing duration in the first and last epochs for *n* = 16 neurons in PR birds where the recording lasted at least 400 s. The average change in duration was*−* 0.03 *±*0.13 s (LMM: *t*_72.5_ =*−* 0.257, *p* = 0.80). (**J**) Comparison of frequency-current slope in first and last epochs for the same neurons. The average change in slope was *−*0.057 *±* 0.027 Hz/pA (*t*_76_ = *−*2.19, *p* = 0.031).

This result rules out the possibility that changes in firing patterns are a consequence of degrading recordings, which can lead to reduced excitability as cells become unhealthy and electrochemical gradients run down. It is unlikely that the neurons in PR birds are inherently healthier than in CR birds, so the absence of plasticity in PR animals implies a specific mechanism that is only active in birds that have been exposed to the colony environment.

### Intrinsic plasticity requires potassium currents

To further confirm whether neurons in CR birds were becoming more phasic because of Kv1.1, we tested whether the plasticity could be reversed by blocking low-threshold potassium currents. We showed previously that application of 4-AP causes phasic cells to become tonic (Chen and Meliza, 2018); here, we recorded from tonic neurons until they became phasic, and then applied 4-AP in the bath (Fig. 5A). As illustrated in Figure 5, this treatment caused neurons to resume tonic firing (Fig. 5B), accompanied by a reduction in outward rectification (Fig. 5C) and in the rheobase (Fig. 5D). All of the plastic neurons fully reverted to tonic firing, while the one stable neuron was unaffected (Fig. 5E). 4-AP did not significantly reverse the decrease in the slope of the f-I curve (Fig. 5F), which is consistent with the suggestion that a Kv1.1-independent process is also contributing to changes in intrinsic properties.

**Fig. 5.**
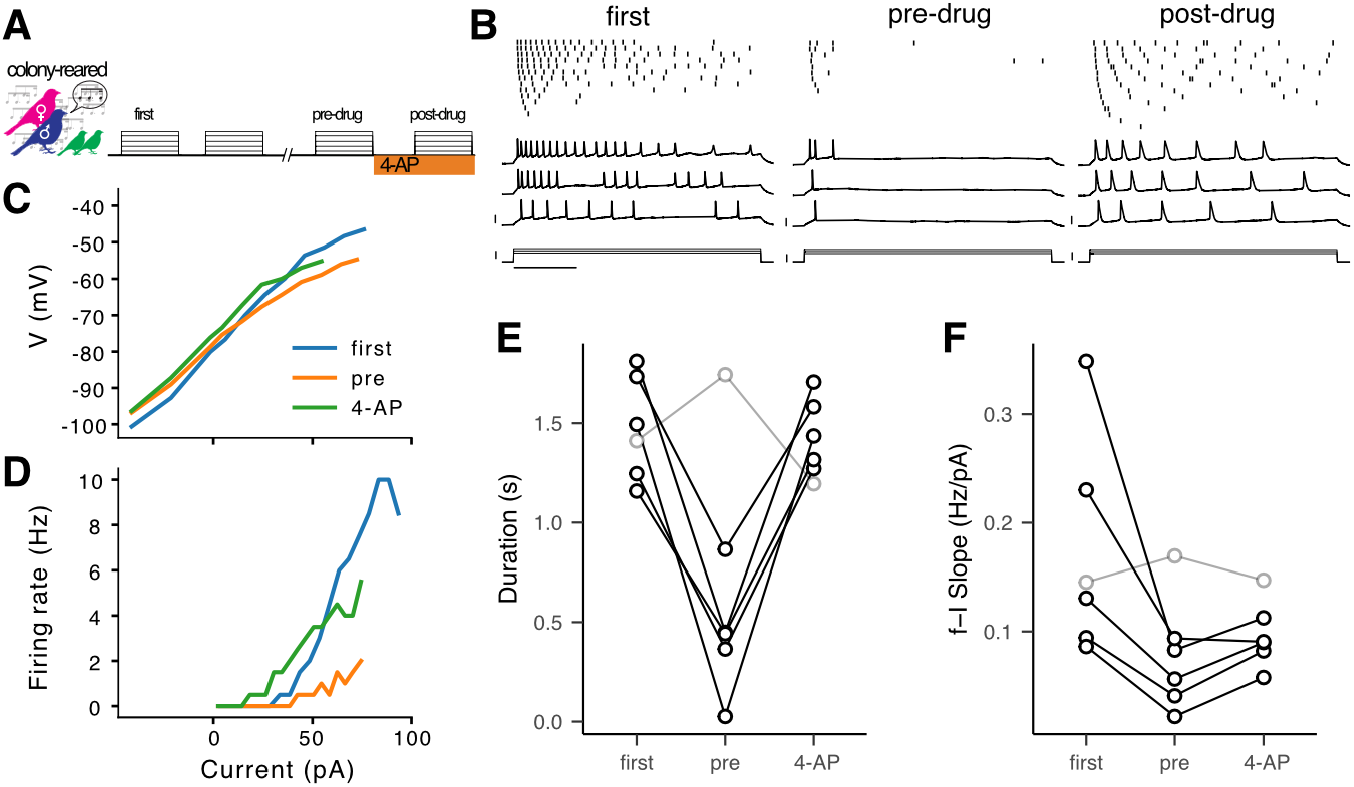
4-AP reverses the shift to phasic dynamics. (**A**) Neurons from CR birds were stimulated with epochs of depolarizing step currents. After 700–1700 s of recording, 4-AP was applied in the bath. (**B**) Responses of an exemplar neuron during the first epoch, the last epoch before drug application (pre-drug), and the epoch after drug application (4-AP). Format is the same as in 3B,C. Scale bars: 40 mV, 50 pA, 500 ms. (**C**) Steady-state voltage versus injected current for first, pre-drug, and post-drug epochs. (**D**) Firing rate versus injected current amplitude for first, pre-drug, and post-drug epochs. (**E**) Comparison of average firing duration in the first, pre-drug, and post-drug epochs for *n* = 6 neurons. One neuron (shown in gray) remained tonic and was not further analyzed. For the remaining neurons, the average change in duration from first to pre-drug was *−*1.00 *±*0.14 s (LMM: *t*_156_ = 7.7, *p <* 0.001), and the change from pre-drug to post-drug was 0.96*±* 0.12 s (*t*_155_ = 7.8, *p <* 0.001). There was no significant difference in duration between the first epoch and the post-drug epoch (*t*_154_ = − 0.29, *p* = 0.95). (**F**). Comparison of frequency-current slope in the same neurons. 4-AP did not significantly reverse the change in slope (mean change = 0.028 *±* 0.034 Hz/pA, *t*_8_ = 0.81, *p* = 0.44).

### Intrinsic plasticity depends on activity and intracellular calcium

Neuronal activity drives an increase in Kv1 currents in CA1 cells of the mammalian hippocampus (Morgan et al., 2019) and in the avian nucleus magnocellularis (Adachi et al., 2019), so we hypothesized that intrinsic plasticity in CM was also triggered by activity. It was not possible to completely eliminate neural activity, because we had to evoke spiking in order to measure neuronal firing patterns. Instead, we used a minimal stimulation protocol with only a single epoch recorded at the beginning and end of the experiment, and the neuron was voltage-clamped at its resting potential in the interim (Fig. 6A). As predicted, neurons were much less plastic when they were stimulated less (Fig. 6B). Only one of the cells (6%, *n* = 1/15) showed a decrease in average duration of more than 1 s, and there was no significant change in duration or f-I slope.

**Fig. 6.**
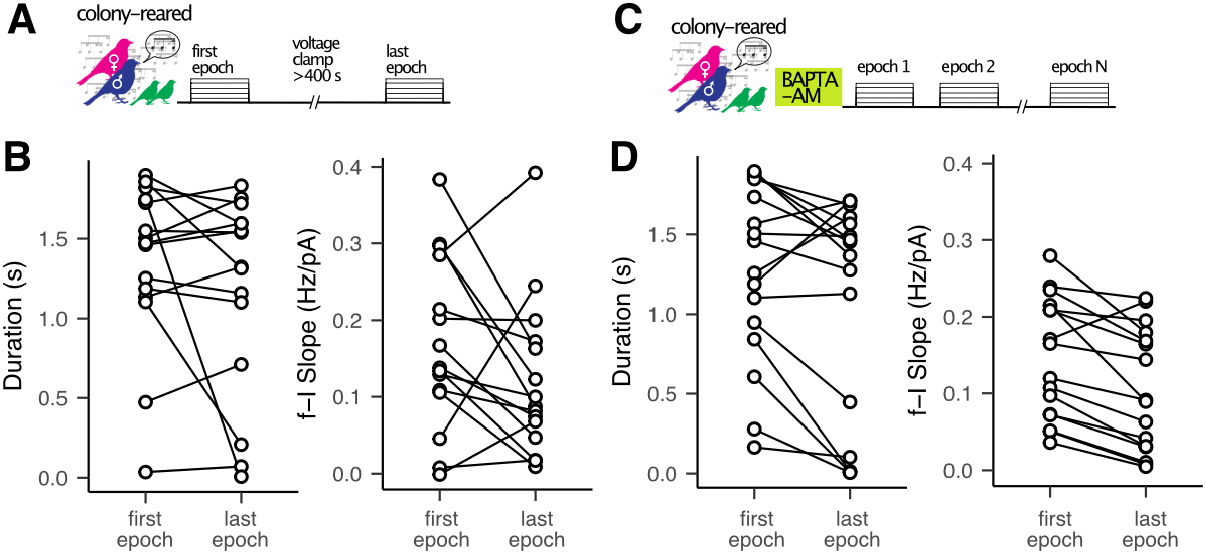
Intrinsic plasticity depends on activity and intracellular calcium (**A**) In the minimal stimulation condition, neurons from CR chicks were tested with a single epoch of depolarizing step currents, clamped at the resting potential for at least 400 s, and then tested with another epoch of current steps. (**B**) Comparison of average firing duration (left) and f-I slope (right) in the first and last epochs for the minimal stimulation condition (*n* = 15). The average change in duration was *−*0.18 *±*0.13 s (LMM: *t*_66.2_ = *−*1.38, *p* = 0.17), and the average change in slope was *−*0.048*±* 0.027 Hz/pA (*t*_76_ =*−* 1.79, *p* = 0.078). (**C**) In the BAPTA condition, slices from CR chicks were pre-incubated with 10 *µ*M BAPTA-AM for 30 min. Cells were stimulated with epochs of depolarizing step currents for at least 400 s. (**D**). (**B**) Comparison of average firing duration (left) and f-I slope (right) in the first and last epochs for the BAPTA condition (*n* = 16). The average change in duration was *−*0.18 *±* 0.13 s (*t*_71.4_ = *−*1.37, *p* = 0.17), and the average change in slope was *−*0.041 *±* 0.026 Hz/pA (*t*_76_ = *−*1.60, *p* = 0.11).

One possibility whereby activity could trigger export of Kv1.1 to the plasma membrane is through an increase in intracellular calcium concentrations. To test this hypothesis, intracellular calcium dynamics were clamped using the fast calcium chelator BAPTA, administered by pre-incubating the slices with the AM ester of BAPTA (Fig. 6C). With this treatment, there was no significant change in duration or f-I slope(Fig. 6D).

A significant change in firing duration was only observed in CR birds under normal recording conditions. To compare plasticity across all the conditions tested (and avoid the fallacy that a difference in significance implies a significant difference), the data were analyzed in unified linear mixed-effects models, one for firing patterns (Fig. 7A) and one for excitability (Fig. 7B). On average, spiking duration decreased by over 500 ms in CR neurons under normal conditions, significantly more than in PR neurons, in the minimal stimulation condition, and with BAPTA treatment. The PR, minimal stimulation, and BAPTA conditions were not statistically distinguishable from each other. For excitability (slope of the f-I relationship), there was no significant difference among the conditions. This result shows that there was both a non-specific decrease in excitability across all our experiments and a specific change in firing patterns related to experience, activity, and intracellular calcium.

**Fig. 7.**
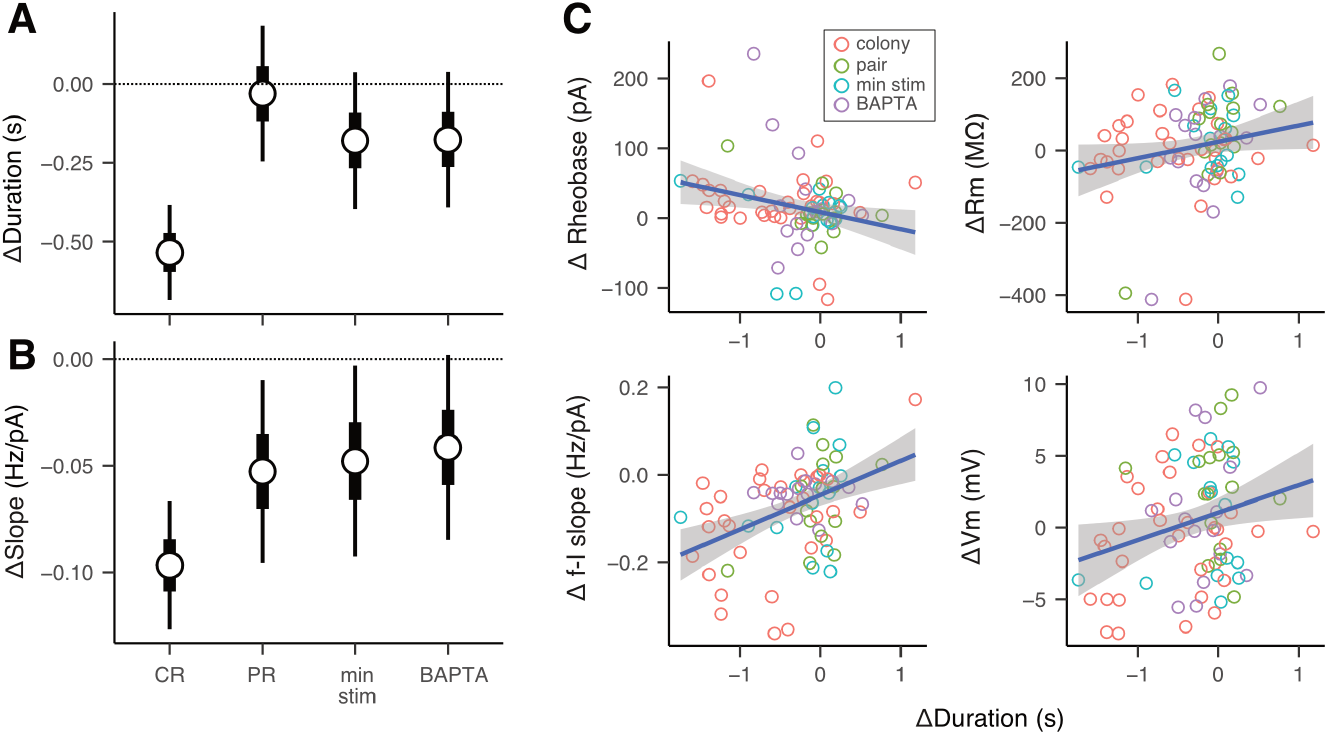
Intrinsic plasticity depends on experience, activity, and intracellular calcium, and is correlated with changes in passive membrane properties. (**A**) Average change in response duration for the CR, PR, minimal stimulation, and BAPTA conditions. Hollow circles show means; thick and thin whiskers show 50% and 90% credible intervals for the mean. The change in duration in the CR condition is greater than for the PR condition (*t*_73.2_ = 3.25, *p* = 0.002), the minimal stimulation condition (*t*_68.8_ = 2.32, *p* = 0.023), and the BAPTA condition (*t*_72.4_ = 2.35, *p* = 0.021). The differences between PR, minimal stimulation, and BAPTA conditions were not significant (*p >* 0.42 for all comparisons) (**B**). Average change in f-I slope for the same conditions. The differences were not statistically significant (*F*_3,76_ = 1.62, *p* = 0.19). (**C**) Comparison of changes in duration with changes in rheobase, input resistance (Rm), f-I slope, and resting membrane potential (Vm). Each symbol represents a single neuron, with color corresponding to the experimental condition. The blue line is the best linear fit, and the gray band is the standard error of the fit. Across all conditions, a decrease in duration was correlated with an increase in rheobase (*r*_78_ = *−* 0.27, *p* = 0.017), an increase in f-I slope (*r*_78_ = 0.42, *p <* 0.001), and a decrease in Vm (*r*_78_ = 0.26, *p* = 0.021). Changes in duration were not significantly correlated with changes in Rm (*r*_78_ = 0.22, *p* = 0.054).

In further support of a specific mechanism, the changes in duration we observed were correlated with changes in intrinsic properties that would be expected from an increase in the conductance of a Kv1.1-like potassium current with an activation voltage close to the resting membrane potential (Fig. 7C): decreasing excitability (f-I slope), decreasing resting membrane potential (Vm), and increasing rheobase. Although the change in input resistance (Rm) was not statistically significant, the trend was also in a direction expected from increased resting conductance (and therefore lower Rm). Changes in duration were not accompanied by what is typically seen during rundown (increasing Vm) or closing of the opening from the recording pipette into the cell (increasing Rm).

## Discussion

These results demonstrate that neurons in the caudal mesopallium exhibit rapid, activity-dependent plasticity of intrinsic dynamics when zebra finches are near the beginning of the sensitive period for song memorization (P30–P35). This plasticity requires early exposure to an acoustically complex environment, which causes elevated expression of the low-threshold potassium channel Kv1.1 in the endoplasmic reticulum and plasma membrane (Fig. 1). Plasticity is primarily observed in tonic-spiking neurons: suprathreshold stimulation for more than a few minutes causes a shift to phasic spiking (Fig. 3) that is reversed by blocking low-threshold potassium currents (Fig. 5) and prevented by blocking intracellular calcium signaling (Fig. 6). In constrast, phasic neurons almost never become tonic. Spiking patterns are not plastic in neurons from pair-reared birds (Fig. 4), which have much lower levels of Kv1.1 expression. Taken together, these findings support a model in which spiking triggers calcium-dependent trafficking of Kv1.1 from intracellular compartments to the plasma membrane. This rapid form of plasticity could facilitate early auditory development by allowing neurons in this higher-order brain region to nimbly adjust their excitability and temporal integration properties in response to the challenging “cocktail-party” acoustic background of the colony.

The present study sheds light on the mechanisms underlying the diverse intrinsic dynamics in CM (Chen and Meliza, 2018) and their dependence on age and experience (Chen and Meliza, 2020). Phasic spiking appears to require both a slow process of Kv1.1 synthesis and a rapid process of Kv1.1 mobilization, as illustrated in Figure 8. The slow process begins around the time zebra finches fledge from the nest and reflects the environment the chicks experience over the next two weeks. Compared to pair-reared chicks, colony-reared chicks experience a louder and more complex acoustic environment, which may result in increased activity-dependent Kv1.1 expression, as seen in the avian auditory hindbrain (Lu et al., 2004; Akter et al., 2018). Non-auditory and neuromodulatory factors could also contribute, as CR birds would be able to engage in more visual and vocal interactions with other animals in the colony. By the time CR birds reach P30–P35, almost all of the excitatory neurons express Kv1.1, but only some of these exhibit phasic spiking at the start of recording, and there is no correlation between intracellular Kv1.1 expression and spiking dynamics (Fig. 2). However, many of the tonic neurons are plastic, with the potential to become phasic when they are stimulated, suggesting that the intracellular Kv1.1 is a ready-to-use pool that can be rapidly mobilized to the plasma membrane to alter intrinsic dynamics.

**Fig. 8.**
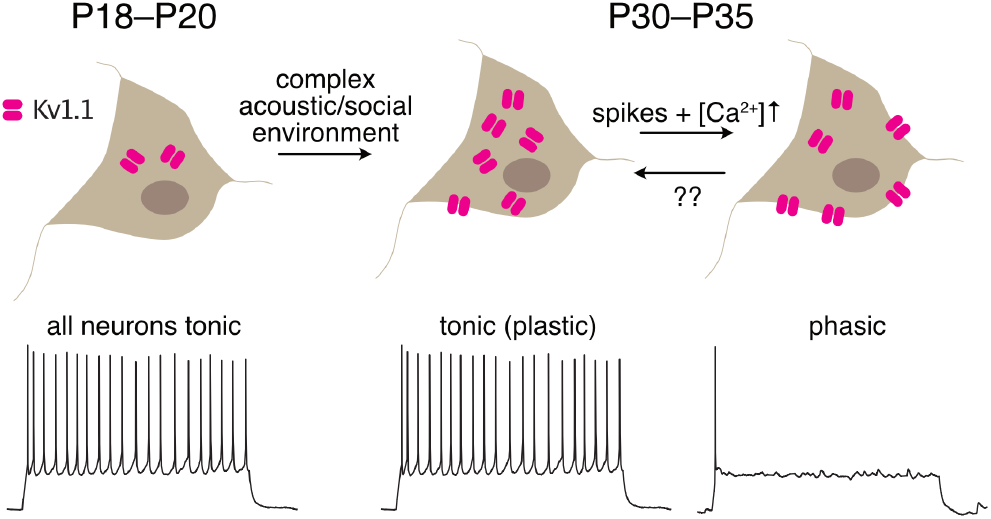
Model of slow and fast intrinsic plasticity. When zebra finches fledge (P18–P20), CM neurons have low levels of Kv1.1 expression and are essentially all tonic. Exposure to a complex environment stimulates Kv1.1 expression in CR birds, but neurons remain tonic so long as the Kv1.1 remains within the neuron. Acute stimulation that elevates intracellular calcium causes Kv1.1 to be exported to the plasma membrane, rapidly shifting the dynamics to phasic spiking. Kv1.1 expression remains low in PR animals, so their neurons are not able to become phasic. An unknown process may allow phasic neurons to internalize Kv1.1 and shift back to tonic spiking.

Although changes in intrinsic dynamics can be the consequence of non-specific rundown, all of the evidence points to a specific mechanism involving a low-threshold potassium current that is likely mediated at least in part by Kv1.1. First, we did not observe plasticity in PR birds, which have much lower levels of Kv1.1 expression. Second, plasticity was reversed by blocking low-threshold potassium channels with 4-AP. Third, plasticity required a minimum level of stimulation, typically occurring only after at least 120 spikes. Finally, changes in duration were not associated with changes in passive membrane properties that typically accompany degraded recordings. Further evidence could be obtained with more direct measures of Kv1.1 trafficking and low-threshold potassium currents (e.g., with live microscopy or voltage-clamp recordings), but these experiments would likely require using dissociated cell culture to obtain better optical and electrical access.

The time course of the plasticity and the abundance of Kv1.1 in the ER suggest that Kv1.1 is being mobilized by trafficking to the plasma membrane. Kv1.1 is retained in the ER by binding of the external pore region of the channel to matrix metalloprotease 23 (Rangaraju et al., 2010; Vacher and Trimmer, 2012), which can be cleaved by the calcium-dependent endoprotease furin (Galea et al., 2014). As proposed by Adachi et al. (2019), activity-dependent elevation of intracellular calcium could thereby trigger release Kv1.1 from the ER to the plasma membrane. This model is consistent with our observation that pretreating neurons with the calcium chelator BAPTA-AM blocks plasticity (Fig. 6). Intracellular calcium could also regulate Kv1.1 by modifying the phosphorylation state of channels already localized to the membrane, although this explanation is less likely because calcium-dependent kinases typically downregulate Kv1 conductance (Hoffman and Johnston, 1998) or have only long-lasting effects on expression (Winklhofer et al., 2003).

Activity-dependent intrinsic plasticity has been observed widely in the brain, involving a diverse range of ion channels and mechanisms for modulating their conductances (Zhang and Linden, 2003; Titley et al., 2017; Debanne et al., 2019; Daou and Margoliash, 2021). Plasticity of the low-threshold (D-type) potassium current mediated by Kv1.1 and other members of the Kv1 family has been reported in the cortex (Dehorter et al., 2015; Gainey et al., 2018) and avian nucleus magnocellularis (Lu et al., 2004; Adachi et al., 2019), and also in the hippocampus, where it can occur on the fast time scales we see in CM (Morgan et al., 2019). Whereas hippocampal neurons require intense stimulation to trigger intrinsic plasticity, CM neurons are much more sensitive, requiring as few as 100 spikes over 400 s. Given that we almost never observed phasic cells becoming tonic, this raises the question of why only about half of the neurons in CM are phasic at this age in CR birds. An equilibrium of 50% phasic neurons would require equal rates of tonic cells becoming phasic and phasic cells becoming tonic. Moreover, phasic neurons are essentially absent in CM by P65 (Chen and Meliza, 2020). Therefore, it is likely there is another mechanism of plasticity that can induce phasic-spiking neurons to become more tonic. This putative mechanism could involve sustained inhibitory or metabotropic inputs (Cudmore et al., 2010; Campanac et al., 2013) linked to the closing of the critical period for sensory learning (London, 2017) or more general maturation of the auditory cortex. It is also possible that the tonic, plastic neurons we recorded were comparatively silent *in vivo* up until the time when we cut slices. This could occur if these neurons were not yet strongly integrated into the developing circuitry of CM and the broader auditory system, and indeed intrinsic plasticity may be an essential mechanism for maintaining network homeostasis as connectivity develops. Given the critical importance of this age to the development of vocal production and perception in these animals, it is perhaps unsurprising that neurons in the auditory cortex are able to so rapidly and sensitively change their intrinsic dynamics.

## Acknowledgements

This work was supported by National Institutes of Health grant 1R01-DC018621, National Science Foundation grant IOS-1942480, and the University of Virginia Brain Institute.

